# A transdiagnostic study of children with problems of attention, learning and memory (CALM)

**DOI:** 10.1101/303826

**Authors:** Joni Holmes, Annie Bryant, the CALM Team, Susan E. Gathercole

**Affiliations:** MRC Cognition and Brain Sciences Unit, University of Cambridge, 15 Chaucer Road, Cambridge CB2 7EF, England

## Abstract

**Background:** A substantial proportion of the school-age population experience cognitive-related learning difficulties. Not all children who struggle at school receive a diagnosis, yet their problems are sufficient to warrant additional support. Understanding the causes of learning difficulties is the key to developing effective prevention and intervention strategies for struggling learners. The aim of this project is to apply a transdiagnostic approach to children with cognitive developmental difficulties related to learning to discover the underpinning mechanisms of learning problems.

**Methods / Design:** A cohort of 1000 children aged 5 to 18 years is being recruited. The sample consists of 800 children with problems in attention, learning and / memory, as identified by a health or educational professional, and 200 typically-developing children recruited from the same schools as those with difficulties. All children are completing assessments of cognition, including tests of phonological processing, short-term and working memory, attention, executive function and processing speed. Their parents/ carers are completing questionnaires about the child’s family history, communication skills, mental health and behaviour. Children are invited for an optional MRI brain scan and are asked to provide an optional DNA sample (saliva).

Hypothesis-free data-driven methods will be used to identify the cognitive, behavioural and neural dimensions of learning difficulties. Machine-learning approaches will be used to map the multi-dimensional space of the cognitive, neural and behavioural measures to identify clusters of children with shared profiles. Finally, group comparisons will be used to test theories of development and disorder.

**Discussion:** Our multi-systems approach to identifying the causes of learning difficulties in a heterogeneous sample of struggling learners provides a novel way to enhance our understanding of the common and complex needs of the majority of children who struggle at school. Our broad recruitment criteria targeting all children with cognitive learning problems, irrespective of diagnoses and comorbidities, are novel and make our sample unique. Our dataset will also provide a valuable resource of genetic, imaging and cognitive developmental data for the scientific community.

## Background

Up to 15% of the school population are recognised as having special educational needs (Department for Education, 2017). This group have problems that vary from difficulties in mastering language, reading and mathematics through to attention deficit hyperactivity disorder (ADHD), and many children have multiple areas of difficulty. For most children who are struggling academically, additional support is provided through education services within the school setting. Others also receive specialist interventions through health services including CAMHS (for ADHD) and speech and language therapy services. The long-term economic and social outcomes of this common and highly heterogeneous group of struggling learners include low rates of employment (de Beer, Engels, Heerkens & van der Klink, 2014; Parsons & Bynner, 2005; Emerson & Hatton, 2008; Whitehurst & Lonigan, 1998) and increased risks of mental health and behavioural problems (Emerson & Hatton, 2007). Understanding the underlying causes of these problems provides the key to advancing the development of targeted intervention and prevention strategies and ameliorating these adverse outcomes.

The current study adopts a transdiagnostic approach to identifying the cognitive, behavioural, neural and genetic mechanisms underpinning learning difficulties. It moves away from investigating tightly-defined deficits related to highly specific developmental impairments of cognition towards studying multiple levels the mechanisms and dimensions of disorder in a heterogeneous population. This approach is strongly endorsed by the RDoC NIMH project, in which the primary focus to date has been on psychiatric conditions including mood disorders and psychoses (Cuthbert & Insel, 2013; Doherty & Owen, 2014). It is now widely recognised as equally valuable for cognitive developmental disorders in which there are also high levels of comorbidity, high variability in symptoms for individuals with specific diagnoses and high-levels of cooccurrence of symptoms across different areas of learning difficulty (Casey, Oliveri, & Insel, 2014; Sonuga-Barke & Coghill, 2014; Zhao & Castellanos, 2016). In putting aside singular diagnostic categories, the aim is to understand and characterise the (possibly multiple) dimensions of disorder at the level of the individual child, guiding effective choice of intervention.

Levels of comorbidity across different aspects of learning difficulties are high. Reading difficulties are estimated to co-occur up to 50% of the time with maths (Moll, Kunze, Neugoff, Bruder & Schulte-Korne, 2014) or language problems (McArthur, Hogben, Edwards, Heath & Mengler, 2000). Symptom variability is high within disorders (e.g. Castellanos, Sonuga-Barke, Scheres, Di Martino, Hyde & Waters, 2005) and common cognitive deficits (for example, in phonological skills, working memory (WM), and executive functions (EFs) extend across disorders of reading, maths and language (e.g. Bishop & Snowling, 2004; Wang & Gathercole, 2013; Moll, Göbel, Gooch, Landerl, & Snowling, 2016; Ramus, Marshall, Rosen, & van der Lely, 2013; Szucs, Deine, Soltesz, Nobes & Gabriel, 2013).

The aim of this study is to apply a transdiagnostic approach to children with cognitive developmental disorders related to learning, with the aim of discovering the underpinning mechanisms of disorder. The plan is to recruit a broad sample of children with problems of attention, learning and/or memory (CALM, n=800) and a school-matched group of children who are developing typically (TD, n=200). Recruitment of the CALM group began in 2014 and will be completed in the summer of 2018. These children have been recruited through health and education professionals supporting children who meet the inclusion criteria. Formal diagnoses are not required and no exclusions are made on the basis of comorbid psychiatric, psychological or physical health conditions. Exclusionary criteria are non-native English speakers, uncorrected sensory impairments and the confirmed presence of genetic or neurological conditions known to affect cognition. Recruitment of the TD group will be via schools attended by multiple children in the CALM group and will commence in autumn 2018.

All children complete a broad set of assessments of cognitive abilities known to be impaired in children with learning difficulties including tests of phonological processing, STM and working memory, executive function, attention and fluid reasoning (IQ). They are also given a set of learning measures assessing maths, language and literacy skills. At the time of the clinic visit, children are offered an optional MRI brain scan and asked to provide an optional saliva DNA sample. Parents / carers complete multiple questionnaires about family history and the child’s behaviour, mental health and communication skills. The breath of the recruitment criteria, the scale of the study and the multiple levels of assessment across behaviour, cognition, the brain, and genes make this study a unique resource for understanding the mechanisms of learning difficulties in childhood. The dataset will be made open to the scientific community within 6 months of the completion of data collection and cleaning. We anticipate that this will be in 2020.

## Aims and hypotheses

The primary aim is to use data-driven, hypothesis-free methods to identify dimensions that characterise children based on cognition, behaviour and brain. Adopting a systems neuroscience approach, we will map between these different levels of explanation. Secondary aims are to define groups of children with common cognitive, neural and behavioural profiles and mapping dimensions and data-defined groups against traditional diagnostic categories.

DNA samples will allow us to extend the dimensional analyses to the genetic level. This will be achieved primarily through participation in genetic consortia combining genotype data from developmental cohorts for genome-wide screening of speech, language and reading skills. Existing gene expression data (http://www.brain-map.org/) will be combined with neural data from the CALM sample to identify broad gene groups whose regional expression profile matches important brain organizational features within the sample. These will be used to derive polygenic risk scores to explore how underlying genetic mechanisms might relate to differences in brain organization and in turn be associated with specific patterns of cognitive impairment.

Although the primary statistical approach to be adopted in the study is hypothesis-free, the dataset will provide rich opportunities to test theories of development and disorder, as the following two examples show. First, the large sample of children at educational risk provide high levels of power that can be used to tease apart the cognitive pathways that contribute to different aspects of academic learning. For example, the data can distinguish whether working memory plays a unique role in supporting learning (Gathercole & Alloway, 2008; Gathercole, Alloway, Kirkwood, Elliott, Holmes & Hilton, 2008; Swanson & Sasche-Lee, 2001) or instead that its links with academic achievement are mediated by core domain-specific skills (Cain, Oakhill & Bryant, 2004; Nation, Adams, Bowyer-Crane & Snowling, 1999; Szucs et al., 2013). Second, data collected from the CALM group include substantial numbers of children both with and without ADHD who have learning difficulties. This will enable us to test whether in the children with ADHD, the learning problems have the same cognitive origins as the children with no ADHD or are at least in part are the disruptive consequences of the hyperactive and impulsive behavior distinguishing this group (Sonuga-Barke, 2002; McGrath et al., 2011).

## Methods and design

### Approval

Ethical approval was granted by the National Health Service (NHS) Health Research Authority NRES Committee East of England, REC approval reference 13/EE/0157, IRAS 127675.

### Design

This is a cohort study collecting individual differences measures of cognition and behaviour alongside MRI and DNA data.

### Recruitment and Procedure

Two groups of children aged 5 to 18 years are being recruited. The CALM group (n=800) are referred via health and education practitioners. These include school Special Educational Needs Coordinators (SENCos), paediatricians, speech and language therapists (SaLTs), or psychiatrists and psychologists working in Child and Adolescent Mental Health Services (CAMHS). The majority of referrers work in the South East of England. Referrers are asked to pass an information pack to families with children who they judge in their professional opinion to have problems in the areas of attention, learning and / or memory. Families send an expression of interest form to CALM if they would like to participate in the study. The research team then contacts the referrer to discuss the child’s problems and asks the referrer to describe the child’s primary reason for referral from a choice of attention, literacy, maths, language, memory problems or general poor educational progress. If the child meets the inclusion criteria a CALM clinic appointment letter is sent to the family. Table 1 shows the likely referral profile for n=800 based on the first n=650 children attending the clinic.

**Table 1.** Number of children by referral route and primary reason for referral (n female) for first 650 children attending CALM

| Category | Attention problems | Literacy problems | Maths problems | Language difficulties | Poor educational progress | Memory problems | Total |
| --- | --- | --- | --- | --- | --- | --- | --- |
| Education | 86 (21) | 52 (18) | 12 (4) | 38 (8) | 138 (51) | 57 (29) | 383 (131) |
| CAMHS<sup>1</sup> & Paediatrics | 134 (33) | 5 (1) | 4 (2) | 19 (3) | 58 (15) | 5 (1) | 225 (55) |
| Speech & language therapy | 3 (0) | 2 (1) | 0 | 18 (8) | 2 (1) | 6 (4) | 31 (14) |
| Total | 223 (54) | 59 (20) | 16 (6) | 75 (19) | 198 (67) | 68 (34) | 639 (200) |

The TD group will be 200 children who are typically developing. They will be recruited from schools attended by 1 or more children in the CALM group. School SENCos who have referred children with difficulties to CALM will provide a point of contact within schools. All children on the school register with exception of those who have already been referred to CALM, those with sensory impairments and those who are non-native English speakers will be invited to participate. Children will be given an information pack in school to take home to their parents / carers, which will contain an expression of interest form to be returned to CALM. Appointments for assessments at the CALM clinic will be made upon receipt of expression of interest forms.

All families attend the CALM clinic at the MRC Cognition and Brain Sciences Unit, University of Cambridge, U.K., for the cognitive and behavioural assessments. At the beginning of the session written consent is obtained from the parent/ carer and verbal assent is taken for the child. The assessment takes approximately 3.5 hours. Families are instructed to administer medication as normal if their child has a prescription, and wear glasses / hearing aids as normal if necessary. Cognitive and learning tasks, plus the child questionnaires, take place one-to-one between the examiner and the child in a dedicated testing room. Families sit in a waiting room outside the testing room and are asked to complete behaviour, family history and mental health questionnaires about the child. For younger children sticker charts are used to motivate the child during the session. All children are awarded a small prize at the end of the session and families are reimbursed for their time and travel.

The assessment protocol has two scheduled breaks. During the first, the child is invited to provide an optional DNA (saliva) sample. Families are asked to provide separate consent and assent for providing optional DNA samples. The child’s height and weight is also measured in this break. During the second break the family is given the opportunity to try a mock MRI scanner. The researcher explains how an MRI scan works and gives the child the opportunity to practice going inside and laying still the mock scanner. At the end of the cognitive testing session, families are invited for an additional visit for the child to have an optional MRI scan. Expressions of interest for scanning are taken at this time and followed up with a telephone call to make a separate appointment and ensure the child is suitable for scanning. Consent and assent for scanning are obtained prior to the MRI scan. All families are asked to provide optional consent to be contacted regarding future research projects.

Families are reimbursed for their time and travel. Following the cognitive and behavioural assessment a report summarising the child’s strengths and weaknesses is sent to referrers of children in the CALM group (n=800) to be used by the referrer to guide their ongoing support for the child.

#### Recruitment Phases

The children (N = 1000) are being recruited in four phases. Diagnostic information supplied by referrers for children recruited in each Phase up to n=650 is provided in Table 2. A CONSORT flow diagram summarising recruitment up to n = 650 is provided in Figure 1.

**Figure 1.**
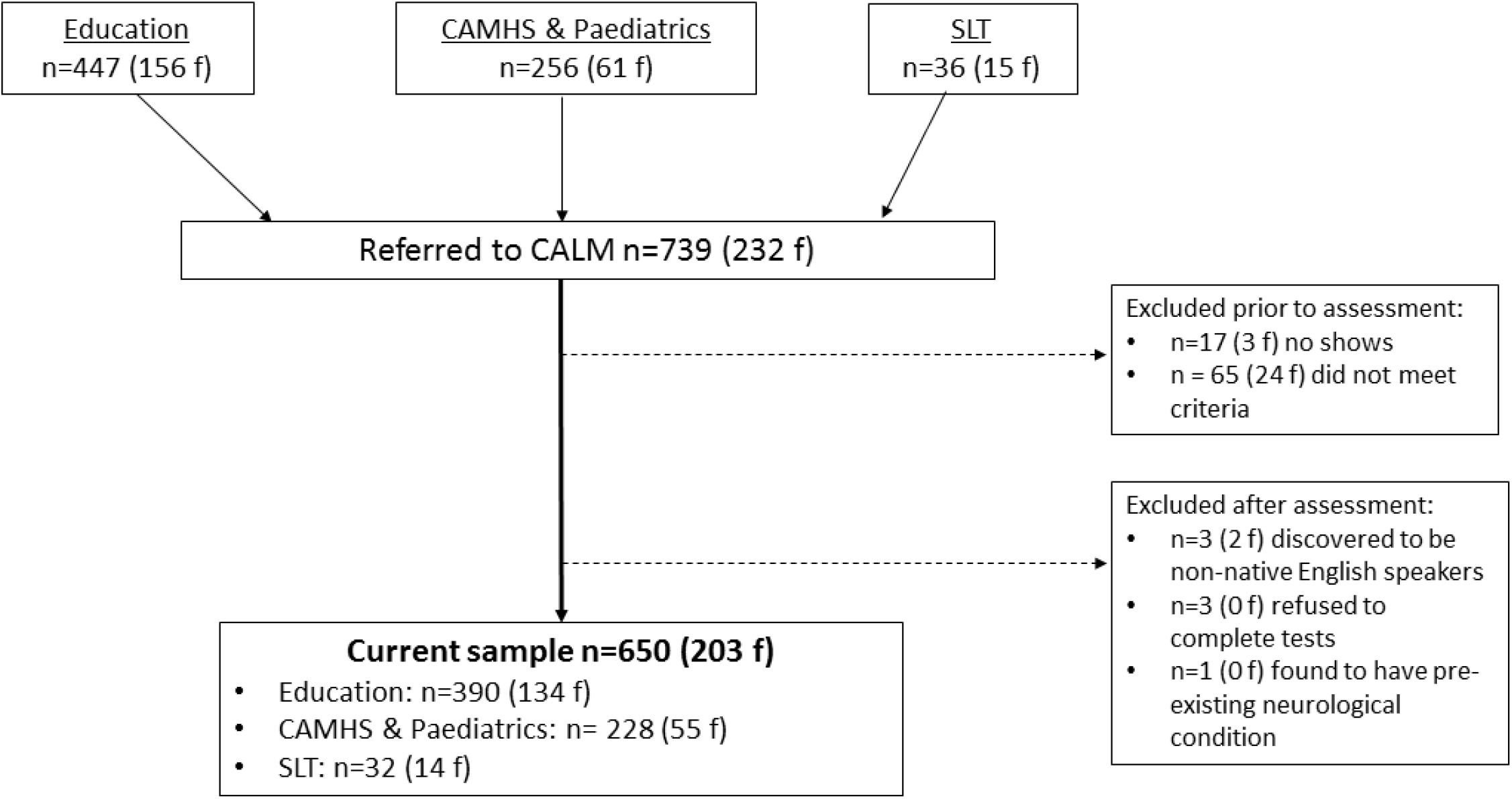
CONSORT flow diagram for first 650 children in the CALM sample

**Table 2.** Diagnostic status of children referred in phases one, two and three for first 650 children attending CALM (n female)

| Phase | One | Two | Three<br>(ongoing) | Total |
| --- | --- | --- | --- | --- |
| ADD | 5 (3) | 6 (3) | 0 | 11 (6) |
| ADHD | 24 (4) | 83 (11) | 30 (10) | 137 (25) |
| Possible ADHD | 5 (1) | 40 (13) | 10 (3) | 55 (17) |
| Hyperactivity | 1 (0) | 0 | 0 | 1 (0) |
| Dyslexia | 22 (8) | 9 (3) | 4 (1) | 35 (12) |
| Dyspraxia | 10 (4) | 5 (0) | 2 (0) | 17 (4) |
| Dysgraphia | 1 (0) | 0 | 0 | 1 (0) |
| Dyscalculia | 0 | 0 | 1 (1) | 1 (1) |
| FASD | 4 (3) | 1 (0) | 1 (1) | 6 (4) |
| Generalised/global<br>delay | 4 (2) | 3 (1) | 0 | 7 (3) |
| Social anxiety | 1 (1) | 0 | 0 | 1 (1) |
| Depression | 2 (2) | 0 | 1 (1) | 3 (3) |
| Autism | 15 (1) | 19 (2) | 8 (1) | 42 (4) |
| PDA | 0 | 1 (1) | 0 | 1 (1) |
| Tourettes | 2 (0) | 2 (1) | 1 (0) | 5 (1) |
| DAMP | 3 (1) | 1 (0) | 0 | 4 (1) |
| Anxiety | 0 | 3 (0) | 3 (1) | 6 (1) |
| OCD | 1 (1) | 2 (1) | 1 (1) | 4 (3) |
| Sensory<br>processing<br>disorder | 0 | 1 (0) | 0 | 1 (0) |
| Known genetic<br>condition | 1 (0) | 1 (0) | 4 (1) | 6 (1) |
| Language disorder | 0 | 1 (1) | 1 (0) | 1 (1) |
| Conduct disorder | 0 | 1 (0) | 0 | 1 (0) |
| ODD | 0 | 1 (0) | 2 (0) | 3 (0) |
| Epilepsy | 2 (1) | 1 (0) | 1 (0) | 4 (1) |
| Speech & language<br>therapy support | 18 (9) | 91 (23) | 14 (4) | 123 (36) |
| No diagnosis | 242 (87) | 103 (28) | 62 (24) | 407 (139) |

##### Phase 1

Between October 2014 and February 2016 children aged between 5 and 18 years who were considered by a health or educational professional to have one or more difficulties in attention, memory, language, literacy and/or maths were recruited. The number of children assessed during Phase 1 was 322 (113 female).

##### Phase 2

Due to the high number of children recruited in Phase 1 without diagnoses priority for referrals in Phase 2 between March 2016 and August 2017 was given to: i) children with ADHD or probable ADHD, classed as having seen an ADHD nurse practitioner and under assessment for a diagnosis by a clinician; ii) those with speech and language problems, defined as having received support from a speech and language therapist within the last two years, or iii) those who have obsessive compulsive disorder (OCD), are on a waiting list to be assessed for OCD, or are currently receiving therapy for OCD traits. The recruitment age was narrowed to 6-12 years of age. The number of children assessed during Phase 2 was 215 (50 female).

##### Phase 3

Having recruited a large number of children with ADHD and many who were receiving support from SaLTs in Phase 2, the Phase 1 recruitment criteria were reinstated in Phase 3 in September 2017. This phase is continuing to recruit until the total n=800 CALM children across Phases 1, 2, and 3 is reached.

##### Phase 4

From autumn 2018, 200 typically developing children aged 5 to 18 years will be recruited through schools attended by children in the first three phases.

### Recruitment criteria

Inclusion criteria for both groups are aged 5 to 18 years and native English speakers (the first language learned and the main language used in the home). All children with cognitive and / or learning problems, as identified by a professional working with them, are accepted into the CALM group irrespective of diagnosis or comorbidities. Children in the TD group will be accepted if they attend the same school as a child in the CALM group and have not been referred to the CALM clinic. Exclusion criteria for both groups are significant uncorrected problems of hearing or vision, pre-existing neurological conditions for which cognitive difficulties are known possible symptoms, and not being a native English speaker.

### Measures

#### Cognition

##### Phonological Processing

Two subtests from the Phonological Assessment Battery (PhAB), Frederickson, Frith & Reason, 1997) are administered. The Naming Speed subtest assesses speed of phonological production. Children are asked to name aloud five drawings of common objects: ball, hat, door, table, and box. They are then presented with a card showing many of these objects and are asked to name them aloud as quickly and accurately as possible. Children complete two trials (cards) and the total completion time in seconds is combined from both trials to give a naming speed raw score. Scores from children who make more than three uncorrected errors per card are treated with caution. The Alliteration subtest measures the ability to isolate initial sounds of simple words. In a series of trials children are presented with three spoken single syllable words and asked to identify which two begin with the same sound. If the children fail to identify correct answers in the three practice trials a supplementary Alliteration Test with Pictures is administered. There are ten trials. Raw scores are the total number of trials correct. Raw scores from both PhAB subtests are converted to standard scores (M = 100, SD = 15).

The Children’s Test of Nonword Repetition (CNRep, Gathercole & Baddeley, 1996) is also given. This assesses phonological processing and shortterm memory. Forty unfamiliar non-words ranging in syllable length from 1 to 4 syllables are spoken aloud one at a time. The child is asked to repeat each word immediately after presentation. Correct scores are given for non-words pronounced correctly. Raw scores out of a possible total of 40 are recorded. The CNRep test was not administered to the first 300 children attending the CALM clinic.

##### Processing Speed

The Visual Scanning and Motor Speed subtests of the Delis Kaplan Executive Function System (Delis, Kaplan, & Kramer, 2001) are administered. Motor speed involves tracing a dotted line to connect circles as quickly as possible. The visual scanning test requires children to cross out all the number threes on a response page of numbers and letters. Errors and time taken to complete the tasks are recorded, and completion times are converted to scaled scores (M = 10, SD=3).

##### Short-term and Working Memory

Four subtests from the Automated Working Memory Assessment (AWMA, Alloway, 2007) are administered. All are span tasks, with 6 trials at each span length. Tasks automatically progress up a span level if there are four or more correct answers within a block and discontinue following three or more incorrect responses. Trials correct are converted to standard scores for each task (M = 100, SD=15). Digit Recall (verbal STM) involves immediate serial recall of sequences of spoken digits. The maximum list length is nine digits. Backward Digit Recall (verbal WM) follows the same procedure except children attempt to recall the memory items in reverse sequence. Maximum list length is set to seven digits. The Dot Matrix subtest (visuo-spatial STM) requires children to recall the locations of a series of dots presented one at a time in a four by four matrix. Up to nine dots can be presented in a sequence. In Mr X (visuo-spatial WM) the child must first decide whether the two Mr X figures are holding a ball in the same hand as each other. The Mr X figure on the left is upright, while the Mr X on the right can be rotated to one of seven positions. The child is asked to remember the location of the ball held by the Mr X on the right, and after successive displays of pairs of Mr Xs the child attempts serial recall of positions in which the ball was held. This task increases up to a maximum of span length of 7.

Children also complete a Following Instructions task developed by Gathercole et al. (Gathercole, Durling, Evans, Jeffcock & Steon, 2008), in which participants are required to carry out sequences of instructions on an array of props laid out in front of them. The instruction sequences consist of descriptions of actions to be performed on a set of five stationery items (a ruler, an eraser, a pencil, a folder, and a box), in each of three colours (red, yellow, or blue). There are two actions: touch (e.g., touch the red pencil) and pick up (e.g., pick up the yellow ruler). Actions involving touching and picking up are concatenated using the adverb “then” to produce increasingly longer sequences that vary in length but not in lexical complexity. A span-type procedure is employed in which the length of the instruction sequence increases systematically. Each span consists of a block of six trials. Testing starts at one action (e.g., Touch the red ruler), increases by one action per block (e.g., touch the red ruler and then pick up the yellow pencil), and is terminated after three incorrect trials in one block. The object array is in view at all times. Participants listen to the instructions and are restricted from manipulating any of the objects. At the end of the presentation, participants are asked to perform the actions in sequence. Responses are recorded as accurate if all elements of the individual action phrase—action, object, and colour—are correctly recalled in their original serial position in the instruction sequence. The number of correct features (colour), objects (item such as pencil / pen etc) and actions (touch pick up) are also recorded.

##### Episodic Memory

The Stories subtest of the Children’s Memory Scale (Cohen, 1997) is used to assess language skills and episodic memory. The child hears two stories (the pairs of stories presented depend on the age of the child). After each story the child is asked to retell the story in as much detail as possible to provide an index of immediate recall. Following a short delay (carrying out a separate task) the child is asked to retell the two stories again (delayed recall), and then asked yes/no factual questions about each story (delayed recognition). Scores of immediate and delayed verbal recall and delayed recognition are converted to scaled scores (M = 10, SD=3).

##### Executive Function

The Tower and Trail Making subtests of the DKEFS are administered to children aged 8 years and above to measure planning and switching abilities respectively. The Tower Test involves building a tower to match a presented picture using five disks of different sizes arranged on three pegs. The child must build the tower in the fewest number of moves possible and as quickly as possible, moving only one disk at a time and without placing any disk on a smaller disk. There are a total of nine towers to build, with increasing time limits for each trial. The time of the first move, total time taken per trial, total number of rule violations and accuracy are recorded. Total achievement scores are converted to scaled scores (M = 10, SD=3). The Trails subtest has five conditions. The Visual Scanning and Motor Speed conditions are described under “Speed” above. The Letter Sequencing and Number Sequencing subtests require children to connect letters in alphabetical order (A to P) or numbers in ascending order (numbers 1 to 16). The switching condition, Number-Letter Sequencing involves connecting letters and numbers in an alternating ascending sequence (e.g. A-l, B-2, C-3 etc). For each condition, completion times are converted to scaled scores (M=10, SD=3). Note that the DKEFS subtests were not administered to the first 60 children attending the CALM clinic.

The Matrix Reasoning subtest of the Wechsler Abbreviated Scales of Intelligence II (WASI-II, Wechsler 2011) is used as an index of general reasoning. Children are presented with incomplete matrices of images and asked to select an image to complete each matrix from a choice of four options. For children up to the age of 8 there are a possible 24 matrices to complete. For children aged 9 years and older there are a possible total of 30 matrices to complete. The test is discontinued when the child selects three consecutive incorrect responses. Trials correct are converted to T-scores (M = 10, SD=10).

##### Attention

The Test of Everyday Attention for Children 2 (TEA-Ch2, Manly, Anderson, Crawford, George, Underbjerg & Robertson, 2016) is administered. Children younger than 8 years old complete three tasks from the TEA-Ch2 J (Manly et al., 2016). Children aged 8 and above complete the TEA-Ch2 A version (Manly et al., 2016) that includes more difficult adaptations of the same three tasks plus one additional measure of set-switching. The Simple Reaction Time subtest measures attention-based reaction time. Children focus on a square centred on a blank screen and press a key as soon as blue blob appears anywhere on screen. The task lasts six minutes on average and average response time in seconds is scored. Sustained attention is measured using the Vigil (8 years +) and Barking (<8 years) subtests that require children to count in their heads the number of auditory items (bleeps or barks) heard at random intervals over ten trials. The number of trials correct is scored. Visual selective attention is assessed using the Hector Cancellation (8 years+) and Balloon Hunt (<8years) subtests. Both are time-limited cancellation tasks requiring children to cross out as many target items (either balloons or circles) as possible in a visual scene presented on paper. There are six scenes in total for Hector Cancellation and four for Balloon Hunt. Each varies by the number of distractor items. The total number of targets correctly identified across all scenes is recorded. The switching task, Reds, Blues, Bags and Shoes, is administered only to children over the age of 8 years. Children first sort four repeating visual items (red or blue bags and shoes) according to colour (red or blue) or use (worn on the hand or foot). In further trials children must switch between the sorting rules after every five items. The raw score is mean reaction time on switch trials. For a TEACH-2 tasks raw scores are converted to scaled scores.

#### Learning

##### Vocabulary

The Peabody Picture Vocabulary Test (PPVT, Dunn & Dunn, 2009) measures receptive vocabulary. It involves selecting one image from four options that represent a stimulus word. Children complete four practice items before beginning the test at a set of 12 items corresponding to their chronological age. A basal set is established when a child completes all 12 items in set with one or no errors. If the child makes more than one error, previous sets are administered in reverse order until the basal set is established. Subsequent sets of increasing difficulty are administered until the ceiling set is established: eight or more errors in a set of 12 items. Children can either respond verbally by saying the number of the correct image, or they can point. The test is untimed. The raw score is the number of items correct (the last item in the ceiling set minus total number of errors). Raw scores are converted to standard scores (M = 100, SD=15).

##### Spelling, Reading and Maths

The Spelling, Word Reading and Numerical Operations subtests of the Wechsler Individual Achievement Test II (WIAT II, Wechsler, 2006) are administered to assess children’s learning. The Spelling test measures spelling using letter sounds initially, progressing to single words that increase in difficulty. The Word Reading test is a measure of single word reading that starts with identifying letters, moves on to selecting words with similar sounds and then reading words that increase in complexity. Numerical Operations measures the ability to solve numerical problems on paper. Beginning with number identification and counting, it progresses to simple and more complex mathematical problems. None of the tests are timed. Raw scores for all three subtests are converted to standard scores (M = 100, SD=3).

The Maths Fluency subtest of Woodcock Johnson III Test of Achievement (WJ-III, Woodcock, McGrew, & Mather, 2007) was administered to the first 68 children attending the CALM clinic. In this assessment, the child is given several sheets of simple maths calculations and has to respond accurately to as many items as possible in three minutes. It was substituted for the WIAT II Numerical Operations test due to consistently low scores. To make sure these low scores reflected maths ability and were not caused by the time constraint in the WJ-III, the WIAT II subtest was introduced. A small number of children completed both maths assessments and there were no significant differences in performance across the tests (p>.05).

#### Behaviour

##### Conners

The Conners 3-Parent Rating Scale Short Form (Conners, 2008) is used to assess symptoms related to ADHD. Parents / carers rate the frequency over the past month of 45 descriptions of problem behaviours. Scores on these items form six subscales consisting of Inattention, Hyperactivity/ Impulsivity, Learning Problems, Executive Function, Aggression, and Peer Relations. The sum of raw scores on each subscale is converted to a *T*-score (*M*= 50, *SD*= 10).

##### BRIEF

The Behavior Rating Inventory of Executive Function (BRIEF, Gioia, Isquith, Guy, & Kenworthy, 2000) questionnaire is completed by parents / carers. It contains 80 statements of everyday problem behaviours related a range of executive function difficulties that are rated for frequency over the past six months. *T*-scores are derived for eight subscales: Inhibit, Shift, Emotional control, Initiate, Working memory, Planning, Organisation and Monitor. Three composite scores are also derived: Metacognition, Behaviour Regulation and Global Executive Function. All raw scores are converted to T-scores (M = 50, SD 10).

##### CCC-2

The Children’s Communication Checklist, second edition (CCC-2, Bishop, 2003) is used to measure communication skills. This 70-item parent / carer rating questionnaire assesses language structure and form, and verbal and nonverbal pragmatic communication. Scaled scores (M = 10, SD=3) are derived for 10 subscales that form three categories measuring different aspects of language use. The first four scales Speech, Syntax, Semantics and Coherence assess language structure, vocabulary use, and discourse, and are areas of communication typically impaired in children with Specific Language Impairments. The next four scales Inappropriate Initiation, Stereotyped Language, Use of Context and Nonverbal Communication index verbal and nonverbal pragmatic communication skills. The final two scales, Social relations and Interests assess aspects of language behaviour that are usually impaired in Autistic Spectrum Disorders.

#### Mental Health

##### Strengths and Difficulties Questionnaire

The Strengths and Difficulties Questionnaire (SDQ, Goodman, 1997) asks the parent/carer to rate 25 items measuring Emotional Symptoms, Conduct Problems, Hyperactivity / Inattention, Peer Relationship Problems and Prosocial Behaviour based on their child’s behaviour in the last six months. The first four subscales are summed to provide a total difficulties score. Age norms are available for all scales with cut-offs for assessing clinical levels of internalising and externalising problems.

##### RCADs

The Revised Children’s Anxiety and Depression Scale (RCADS, Chorpita, Yim, Umemoto & Francis, 2000) and the RCADS – Parent Version (RCADS-P, Chorpita et al., 2000) are questionnaires that measure the frequency of symptoms of anxiety and low mood as rated by the children themselves (RCADS, 25 items) or their parent / carer (RCADS-P, containing 47 items). Total anxiety and total low mood scores are derived for both scales, as is a combined depression and anxiety score. RCADS-P provides subscale scores for separation anxiety, social phobia, generalised anxiety, panic disorder, obsessive compulsive disorder, and major depressive disorder. Raw scores are converted to T-scores for each scale and total scores (M = 50, SD=10). The RCADS questionnaires were not administered to the first 390 families attending CALM. RCADS are scored immediately following the child’s assessment and referrers are informed immediately of scores above the clinically significant cut-offs.

#### Structural MRI

MRI measures are collected in a one-hour session conducted on the same site as the CALM clinic on a 3T Siemens Prisma with a 32-channel quadrature head coil. Prior to scanning, children are introduced to the MRI environment using a realistic mock scanner. All children practice going into the scanner and staying still. To facilitate this, children play an interactive game that teaches them to minimize head movements, which are measured through an accelerometer in a headband.

##### Tl-weighted structural image

A high-resolution 3D T1-weighted structural image is acquired using a Magnetization Prepared Rapid Gradient Echo (MPRAGE) sequence with the following parameters: Repetition Time (TR) =2250 milliseconds; Echo Time (TE) = 3.02 milliseconds; Inversion Time (TI) =900 milliseconds; flip angle =9 degrees; number of slices: 192; voxel dimensions =1mm isotropic; GRAPPA acceleration factor =2; acquisition time of 4 minutes and 32 seconds.

##### T2-weighted structural image

A high-resolution 3D T2-weighted structural image is acquired with a Sampling Perfection with Application optimized Contrasts using different flip angle Evolution (SPACE) with the following parameters: TR = 5060.0 milliseconds, TE =102.9ms; number of slices =29; voxel dimensions =0.6875 mm × 0.6875 mm × 5.2 mm; GRAPPA acceleration factor =2; acquisition time of 1 minutes and 38 seconds.

##### Diffusion-weighted image

Diffusion-Weighted Images (DWI) are acquired with a Diffusion Tensor Imaging (DTI) sequence with 64 diffusion gradient directions with a b-value of 1000 s/mm2, plus one image acquired with a b-value of 0. Other parameters are: TR =8500 milliseconds, TE = 90 milliseconds, voxel dimensions = 2mm isotropic; acquisition time of 10 minutes and 14 seconds.

##### Resting State

To assess brain connectivity at rest, T2*-weighted fMRI data is acquired while participants rest with their eyes closed using a Gradient-Echo Echo-Planar Imaging (EPI) sequence. A total of 270 volumes are acquired, each containing 32 axial slices; TR =2000 milliseconds, TE =30 milliseconds, flip angle = 78 degrees, voxel dimensions = 3 mm isotropic; acquisition time of 9 minutes and 6 seconds.

#### Physiological Measures

##### Saliva DNA

DNA samples are collected from children in vials using the Oragene^®^ DNA selfcollection kits. Children are asked to produce a saliva sample by first rubbing their cheeks gently for 30 seconds to create saliva, and then they are asked to spit in a pot. For children who find it hard to create saliva, a small amount (max ¼ tsp) of white table sugar is available to place on the child’s tongue. The saliva samples are stored in Oragene^®^ kits at room temperature (15-30°C), as per manufacturer instructions until extraction of DNA. DNA is extracted as soon as possible and stored at −80°C at the Wellcome Trust-MRC Institute of Metabolic Science at Addenbrooke’s Hospital.

##### Height and Weight

Children’s height and weight is measured during the first CALM visit. A wall chart is used to measure height in centimetres and a set of floor scales to measure weight in kilograms.

### Statistical analysis

Factor analysis, a statistical method that groups variables based on shared variance, will be used to derive underlying dimensions from the cognitive and behavioural data (e.g. Kotov et al., 2017). This technique has been used to identify dimensions of phonological and non-phonological skills in children with diagnosed SLI and dyslexia (Ramus et al., 2013) and separate latent constructs for inattention and hyperactivity in children with ADHD (Martel, von Eye & Nigg, 2010).

Machine-learning approaches will be used to map the multi-dimensional space of the cognitive measures. These methods have rarely been applied to understanding developmental disorders (e.g. Fair, Bathula, Nikolas & Nigg, 2012) - the only applications involve using supervised machine learning in which the learning algorithm attempts to learn about pre-defined categories of children (Peng, Lin, Zhang & Wang, 2013). An unsupervised machine learning approach will be used to learn about the composition of the sample: how children group together across multiple cognitive domains. These approaches will be combined with ways of grouping children according to common cognitive, neural or behavioural profiles. Such methods will include class-based analyses (e.g. latent class or cluster analyses) and clustering algorithms that have been previously used to identify groups of children with distinct learning profiles (Archibald, Cardy, Joanisse & Ansari, 2013).

Direct group comparisons will be made via MANOVAs to test particular hypotheses as the dataset is formed. Bayesian methods will be employed to evaluate the strength of the evidence for and against the null hypothesis in addition to traditional null hypothesis testing (e.g. Kass & Raftery, 1995).

## Discussion

Supporting adults with learning difficulties costs the UK’s NHS £560 million per year for inpatient care. Local authorities and adult social services spend a further £5.3 billion on community services (National Audit Office, 2015). Using evidence-based approaches to understand and address the causes of learning problems in childhood is the key to delivering social and economic benefits (National Institute for Health and Care Excellence, 2015). Our multi-systems approach to identifying the cognitive, neural and genetic dimensions of children’s learning difficulties provides a novel way to enhance our understanding of the common and complex needs of the majority of children who struggle at school, and in doing so illuminates potential targets for intervention for individuals.

Our approach has several strengths.

- It is a large-scale study designed to identify the dimensional basis of learning disorders that adopts a systems neuroscience approach spanning cognition, behaviour, the brain and genes.
- It identifies dimensions that can be used to inform the development of interventions necessary to meet the needs of the individual child.
- It will recruit a heterogeneous sample of poor learners, irrespective of diagnoses and comorbidities, which is highly representative of the majority of children struggling at school.
- It will include a comparison group of typical learners to quantify the size of impairment(s) in poor learners.
- It will provide a rich source of data for testing theories of cognitive development and disorder.
- It will generate a database of developmental data to be made openly accessible to the scientific community 6 months after study completion.
- The data generated by the project directly address the common and comorbid cognitive developmental difficulties faced within school and in the health services, and the outcomes are of direct relevance to these communities. The CALM project website (http://calm.mrc-cbu.cam.ac.uk/) is designed to promote practitioner-researcher working in these areas and to facilitate knowledge transfer to the international community of interested professional groups.

The study has the following limitations.

- Recruitment is restricted to non-native English speakers due to restricted availability of standardised measures.
- Some areas of assessment were very limited. In particular, direct tests of language function were limited to a receptive measure of vocabulary only.
- The DKEFS tests of executive were restricted to children 8 years and older.
- Some assessments were introduced after recruitment had started, generating complete data. These include the CNRep and RCADS.

In summary this study has the potential to make a significant contribution to our understanding of the causes of common learning problems faced by many children in school. Identifying dimensions that distinguish individuals will provide targets for tailored individual interventions.

## Declarations

### Acknowledgements and funding

The Centre for Attention Learning and Memory (CALM) research clinic is based at and supported by funding from the MRC Cognition and Brain Sciences Unit, University of Cambridge. The Principal Investigators are Joni Holmes (Head of CALM), Susan Gathercole (Chair of CALM Management Committee), Duncan Astle, Tom Manly and Rogier Kievit. Data collection is assisted by a team of researchers and PhD students at the CBSU that includes Annie Bryant, Fánchea Daly, Francesca Woolgar, Sally Butterfield, Joe Bathelt, Erin Hawkins, Sinead O’Brien, Silvana Mareva, Amy Johnson, Cliodhna O’Leary, Joe Rennie, Mengya Zhang, Delia Fuhrmann, Lara Bridge. The authors wish to thank the many professionals working in children’s services in the South-East and East of England for their support, and to the children and their families for giving up their time to visit the clinic.

## Data availability

The data will be made openly accessible to the scientific community 6 months after study completion.

